# Development of Sensorimotor-Visual Connectome Gradient at Birth Predicts Cognitive Outcomes at 2 Years of Age

**DOI:** 10.1101/2023.09.07.556778

**Authors:** Yunman Xia, Yuehua Xu, Dingna Duan, Mingrui Xia, Tina Jeon, Minhui Ouyang, Lina Chalak, Nancy Rollins, Hao Huang, Yong He

## Abstract

Functional connectome gradients represent fundamental brain organizing principles. Here, we investigate the maturation of the macroscopic connectome gradients in a task-free functional MRI dataset consisting of 39 preterm and term babies aged 31-42 postmenstrual weeks and its connection with postnatal cognitive growth. We show that the principal sensorimotor-to-visual gradient is present in baby connectomes around 35.6 postmenstrual weeks and continuously develops towards a more term-like pattern. The gradient’s global topographical structure undergoes progressive maturation with age, characterized by increases in explanation ratio, gradient range and gradient variation. The development of focal gradients is mostly localized in primary regions serving sensory, motor and visual functions. Machine learning approaches revealed that the connectome gradient at birth significantly predicts individual cognitive outcomes at 2-y follow-up. These findings highlight the early emerging rules of functional connectome gradients and have implications for understanding how the connectome gradients at birth underlies cognitive growth in later life.

**Highlights:**

- Sensorimotor-to-visual gradient is present around 35.6 postmenstrual weeks
- Global gradient structure undergoes substantial changes during the third trimester
- Regional gradient develops mostly in primary sensory, motor, and visual regions
- Sensorimotor-to-visual gradient at birth predicts cognitive outcomes at age 2

## Introduction

Human adult brain’s functional connectome exhibits fundamental organizing principles, characterized by two core connectivity gradients, including the principal gradient from the primary sensorimotor and visual cortex to the transmodal regions, and the secondary gradient separating the sensorimotor areas and visual cortex (Margulies et al., 2016, 2022). In recent years, significant research has focused on the primary-to-transmodal gradient in the adult brain connectome, highlighting its crucial implications for individual cognitive function (Bethlehem et al., 2020; Wang et al., 2021) and mental health (D. Dong et al., 2023; M. Xia et al., 2020). However, comparatively less attention has been given to the sensorimotor-to-visual gradient. Studies conducted on developmental populations have emphasized the importance of the sensorimotor-to-visual gradient in the neonatal brain (Larivière et al., 2020), showing that its dominance persists into late childhood (H.-M. Dong et al., 2021; Y. Xia et al., 2022). Nevertheless, the development of sensorimotor-to-visual gradient during the prenatal period and its relationship with postnatal cognitive growth remain understudied.

The third trimester of pregnancy is a critical phase for the early development of cortical microstructures and functional organization in the human brain (Cao et al., 2017; Ouyang et al., 2019; Rakic, 1995; Sidman & Rakic, 1973). During this period, cortical neurons and neural circuits undergo rapid and significant growth, including the strengthening of axons (Kostović & Jovanov-Milosević, 2006), the arborization of dendrites (Sidman & Rakic, 1973), and the formation of synapses (Huttenlocher & Dabholkar, 1997). These microstructural changes primarily occur in the primary sensory and motor regions, which are associated with necessary cognitive functions after birth (Huttenlocher, 1990; Rabinowicz, 1986). Functional connectomic studies have demonstrated prominent small-world properties, modular organization, and the emergence of hub regions primarily in the sensorimotor regions during the third trimester (Cao et al., 2017; van den Heuvel et al., 2015). These findings suggest that the primary sensorimotor regions undergo remarkable development during this period. Simultaneously, these regions serve as an anchor in the sensorimotor-to-visual gradient axis. Therefore, we speculated that the development of the sensorimotor regions during the third trimester could drive the emergence and maturation of the sensorimotor-to-visual gradient and that this gradient development could provide neural substrate in support of cognitive growth in later life.

To test the above hypothesis, we collected task-free functional MRI (tf-fMRI) data in 39 neonates aged 31-42 postmenstrual weeks (PMW) at scanning, followed by cognitive assessment at a 2-y follow-up. Using functional connectome gradient analysis, we first identified principal functional gradients in the neonatal brains and their developments with age during the third trimester. Then, we employed support vector regression analysis to examine whether the functional connectome gradients at birth could predict individual cognitive outcomes at 2 years of age.

## Results

### The Emergence of Sensorimotor-to-visual Gradient at 35.6 Postmenstrual Weeks

To qualitatively delineate the development of the sensorimotor-to-visual gradient, we conducted a cross-participant sliding window analysis to investigate changes in connectome gradients across age windows. We first divided all neonates into 15 overlapping age windows, ranging from 33.8 to 40.2 PMW, in ascending order of age. Each window comprised 10 neonates with a step size of two, meaning that window 1 included neonates 1-10, window 2 included neonates 3-12, and so forth until window 15 included neonates 29-39. We then identified a group-level connectome gradient within each age window based on the averaged functional connectivity matrix. As the first two gradients account for most of the variance of functional connectome (22.49% - 36.06%), the present study focuses on the development of the first two gradients during the third trimester. Upon visually inspecting the first two gradients across age windows, we observed that the dominance of the sensorimotor-to-visual gradient emerged at the age window of 35.6 PMW, which is evident through the first-time maximum separation of sensorimotor and visual regions along the first gradient axis (Fig. 1A, *left*). Before this critical age window, the voxels within the sensorimotor area and visual cortex showed undifferentiated locations, lacking the clear separation observed in the later age windows (Fig. 1A, *left*). The cortical mapping for the first two gradients of each age window further confirmed this significant period (Fig. 1A, *right*). Moreover, the secondary gradient exhibited an anterior-to-posterior pattern across the cortex at approximately the window of 34.57 PMW, characterized by the clustered visual cortex localized at the extremes of the second gradient axes (Fig. 1A, *left*). Additionally, we observed that the ranges of both the sensorimotor-to-visual and the anterior-to-posterior gradients gradually extended after age window of 35.6 PMW (Fig. 1A, *left*).

**Figure 1.**
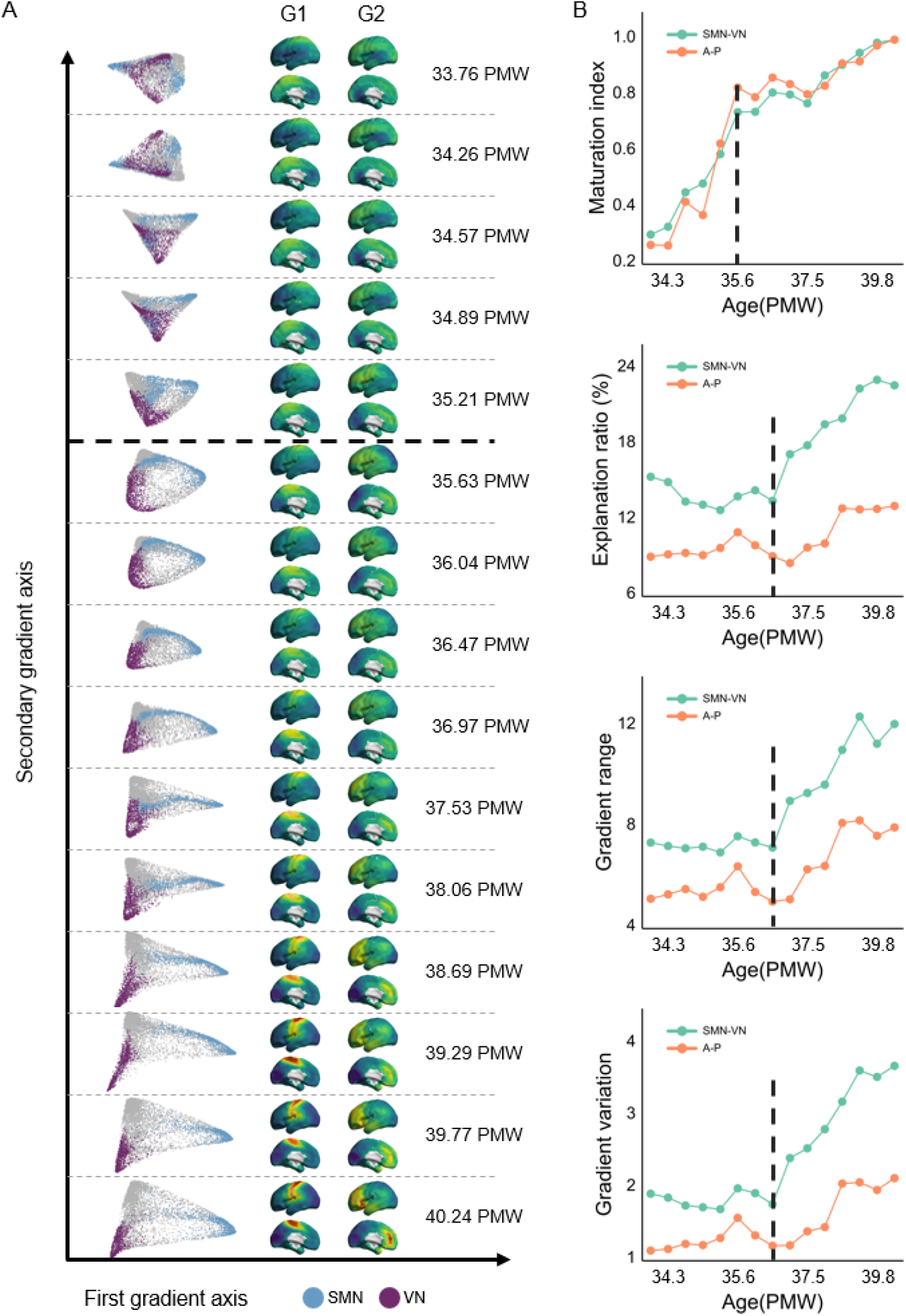
(A) Functional connectome core gradients of 15 age windows, spanning from 33.8 to 40.2 postmenstrual weeks (PMW). In the scatter plots, the *x-axis*, and *y-axis* represent the scores of the first gradient and secondary gradient axis respectively, and the voxels within sensorimotor regions and visual regions were labeled by blue and purple. At the age window of 35.6 PMW, the sensorimotor-to-visual gradient is first observed, indicated by the voxels within the sensorimotor area (*blue*) and visual cortex (*purple*) initially occupying the extreme locations of the first gradient axes. Subsequently, this gradient organization gradually enlarges the ranges along the first gradient axes after this window. The anterior-to-posterior gradient is first observed at the approximate age window of 34.57 PMW, characterized by the voxels within the visual cortex (*purple*) clustered at the extremes of the second gradient axes. G1 means the first gradient, while G2 indicated the secondary gradient. (B) The maturation index and global measures of both gradients across age windows. At the age window of 35.6 PMW, the maturation index of both gradients experiences a significant surge from approximately 0.5 to 0.8, and remains at a high level thereafter. After one week of 35.6 PMW, i.e., the age window of 36.47 PMW, the sensorimotor-to-visual gradient exhibits substantial increases in the explanation ratio, gradient range, and gradient variation, while these metrics remain relatively stable before this age. Moreover, the anterior-to-posterior gradient also demonstrates slight increases in these global measures. SMN-VN means the sensorimotor-to-visual gradient, while A-P indicated the anterior-to-posterior gradient.

To gain insights into the gradient changes across age windows, we calculated the maturation index (i.e., the similarity to the gradients of the oldest age window) and the global measures of core connectome gradients across age windows, including the explanation ratio (i.e., accounting for the variance of the high-dimensional functional connectome), the gradient range (i.e., difference between the values of the positive and negative extremes of the gradient axis), and the gradient variation (i.e., the standard deviation of the values of gradient axis). The results unveiled the maturation indexes of core gradients, i.e., 0.74 in the principle sensorimotor-to-visual gradient and 0.83 in the secondary anterior-to-posterior gradient at 35.6 PMW, which marked a critical time point for the formation of core connectome gradients (Fig. 1B). Interestingly, the sensorimotor-to-visual gradient exhibited substantial increases in the explanation ratio, gradient range, and gradient variation (Fig. 1B) after one week of the emergence of sensorimotor-to-visual gradient (i.e., 36.5 PMW). Conversely, these measures displayed a relatively stable pattern prior to 35.6 PMW (Fig. 1B). Finally, the secondary gradient, i.e., the anterior-to-posterior gradient, demonstrated modest incremental changes across the age windows, as reflected by these three measures (Fig. 1B).

Taken together, these qualitative analyses provide compelling evidence that the sensorimotor-to-visual gradient emerges in neonatal connectomes at around 35.6 postmenstrual weeks, and undergoes continuous development towards a more term-like pattern.

### The Sensorimotor-to-visual Gradient Exhibits Global and Regional Development During the Third Trimester

To quantify the developmental changes in core connectome gradients during the third trimester, we used a general linear model to assess the effect of age on the global measures and regional scores of the first two gradients, with gender and head movement parameters included as covariates.

For the principal sensorimotor-to-visual gradient, we observed significant age-related increases during the third trimester in the maturation index (*t* = 5.02, *P* = 0.0001, Fig. 2A) and all global measures (explanation ratio: *t* = 3.55, *P* = 0.009; gradient range: *t* = 5.67, *P* < 0.0001; gradient variation: *t* = 5.05, *P* = 0.0001, Fig. 2A). For the secondary anterior-to-posterior gradient, we only found significant age-related increase in the maturation index (t =3.85, *P* = 0.004, Fig. 2B). All *P* values were corrected by Bonferroni corrected-*P* < 0.05.

**Figure 2.**
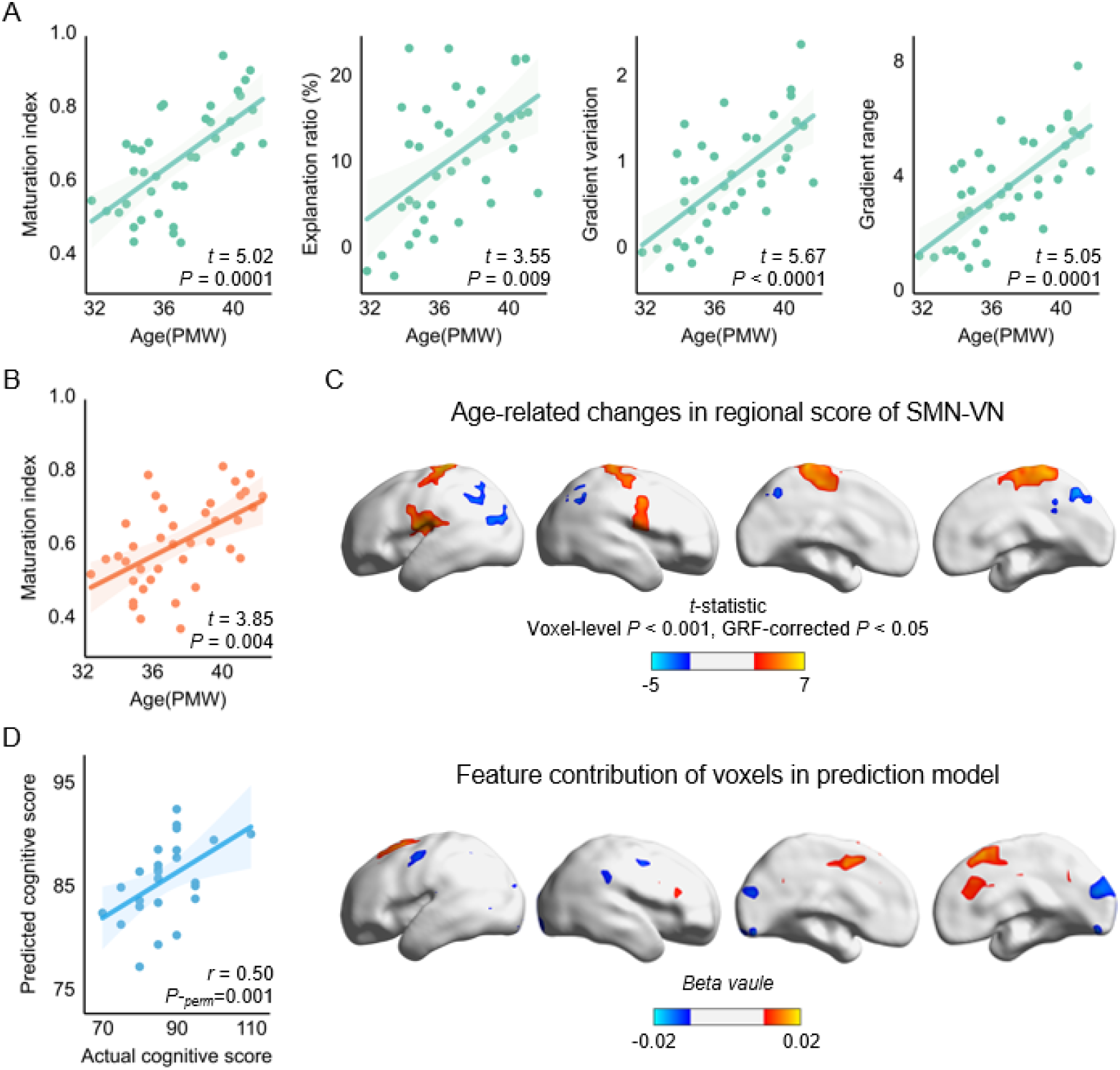
Significant increases with age were seen in (A) the maturation index and all global measures of the sensorimotor-to-visual gradient, and (B) the maturation index of anterior-to-posterior gradient, with gender and head movement parameters included as covariates. Bonferroni-corrected *P* < 0.05. (C) At the regional level, gradient scores that increased with age were mostly found within the sensorimotor regions (shown in warm colors), while gradient scores in the inferior parietal lobe and precuneus decreased with age (shown in cool colors) (voxel-level *P* < 0.001, Gaussian random field (GRF) cluster-level corrected *P* < 0.05). SMN-VN, the sensorimotor-to-visual gradient. (D) The sensorimotor-to-visual gradient from neonates significantly predicts cognitive scores at 2 years of age. The scatter plot shows a significant Pearson correlation between actual cognitive scores and predicted scores (*left*), and the correlation coefficient was corrected by the permutation test (1,000 times). The feature contribution weights of voxels were projected on a cortical surface (*right*). The features with high contribution were mainly located within the sensorimotor and visual cortex.

Voxel-wise statistical analysis revealed age-associated increases in regional gradient scores of sensorimotor-to-visual mainly concentrated in the primary sensorimotor regions, involving the supplementary motor cortex and postcentral gyrus, and age-associated decreases in the inferior parietal lobe and precuneus (voxel-level *P* < 0.001, Gaussian random field cluster-level corrected *P* < 0.05, Fig. 2C). For the anterior-to-posterior gradient, we did not observe significant age effects in the regional scores.

### Sensorimotor-to-visual Gradient at Birth Predicts Cognitive Outcomes at 2 Years of Age

To investigate whether functional connectome gradients in neonatal brains predict individual neurocognitive performance later in life, we employed support vector regression with a 5-fold cross-validation approach. Specifically, we sought to examine whether the regional score of the sensorimotor-to-visual gradient could effectively forecast individual neurocognitive scores at 2 years of age (cognitive, language, and motor scores). During this analysis, we trained models using the whole-brain gradient scores and empirical neurocognitive scores of 21 neonates, leaving out 5 neonates at a time. Subsequently, we utilized these trained models to predict the three neurocognitive scores of each neonate. Notably, among the three neurocognitive scores, we observed a significant association solely between the predicted cognitive component score and the empirical score (*r* = 0.50, *P-perm* = 0.001, permutation test (1,000 times), Fig. 2D, *left*), indicating a reliable prediction of cognitive performance at 2 years of age based on the core connectome gradient in neonatal brains. Moreover, we projected the normalized feature contribution of each voxel onto the cortical surface (Fig. 2D, *right*), revealing that voxels with higher contributions were predominantly situated within the sensorimotor and visual cortex (Fig. 2D, *right*). However, the regional score of sensorimotor-to-visual gradient could not predict the motor and language component scores of individuals at 2 years of age (language: *r* = 0.20, *P-perm* = 0.18; motor: *r*=-0.11, *P-perm* = 0.69). Additionally, the regional score of anterior-to-posterior gradient could not predict all neurocognitive performance at 2 years of age (cognitive: *r* = −0.31, *P-perm* = 0.95; language: *r* = −0.18 *P-perm* = 0.83; motor: *r* = -0.22, *P-perm* = 0.88). Overall, these results underscore the pivotal role of the sensorimotor-to-visual gradient at birth in shaping the postnatal cognitive growth of individuals.

## Discussion

Our study shows for the first-time emergence and development in the functional connectome gradient throughout the third trimester and establishes connections between these gradients in neonatal brains and cognitive performance at two years of age. Specifically, the principle sensorimotor-to-visual gradient showed remarkable changes during the third trimester, including its initial emergence at 35.6 postmenstrual weeks and the subsequent expansion of its global topography, as well as regional variations involving the sensorimotor regions and medial and parietal parietal lobe. Importantly, the sensorimotor-to-visual gradient in neonates significantly predicted individual cognitive scores at 2 years of age. These findings provide valuable insights into the early development of sensorimotor-to-visual gradient, enhancing our understanding of how functional brain organization evolves during pregnancy to support cognitive development in later stages of life.

Although a prior study has demonstrated the existence of the sensorimotor-to-visual gradient in the term-born neonatal brains (Larivière et al., 2020), our study is the first to identify the emergence of the sensorimotor-to-visual gradient at around 35.6 postmenstrual weeks and the age-related changes in this gradient during the third trimester. Notably, previous studies have also observed an abrupt increase in myelinated white matter across the whole brain (Hüppi et al., 1998), and the rapid enhancement in the association between individual variability and the strength of functional connectivity (Xu et al., 2019) at approximately 36 postmenstrual weeks. In conjunction with previous studies, we speculate that the gestational age of 36 weeks is a crucial time point for prenatal development of structural and functional brain organizations, which supports the formation of core gradients of functional connectome. In addition, during the third trimester, the sensorimotor-to-visual gradient shows age-associated increases in the explanation ratio, global topography, and spatial variation, as well as increased gradient scores in the sensorimotor regions. During this period, the sensorimotor regions involved in basic survival functions (Buckner & Krienen, 2013), also undergo remarkable developmental changes. Classical neuroanatomy studies have reproducibly reported the priority maturation in the cytoarchitecture and myeloarchitecture of primary sensory and motor cortices before birth (Huttenlocher, 1990; Huttenlocher & Dabholkar, 1997; Kostović & Jovanov-Milosević, 2006). On the other hand, numerous task-free fMRI studies have indicated that the primary cortex exhibited remarkable increases in the number and strength of functional connectivity, gradually forming an adult-like functional system, as well as serving as the hub nodes of functional brain network (Cao et al., 2017; Eyre et al., 2021; Fransson et al., 2007, 2011). Collectively, the emergence and extension of the sensorimotor-to-visual gradient could be attributable to the gradual maturation of the primary cortex during the third trimester.

In the first 2 years of life, infants gradually develop gross motor and fine motor skills, learn to understand and use words, recognize objects, and form social and self-concepts (Berk, 2015; Rochat et al., 2012). The development of individual mental functions is thought to be a consequence of the maturation of human brain structure and function (Casey et al., 2005; Johnson, 2005). Previous studies suggest that the individual cognitions at two years of age are robustly predicted by neonatal white matter structures and cortical morphological features (Li et al., 2021; Ouyang et al., 2020). The present study highlights that the sensorimotor-to-visual gradient in functional connectome at birth can well predict the cognitive scores at two years of age. These results form a consensus that the establishment of brain functional organization during the third trimester has important implications for postnatal cognitive growth. The sensorimotor-to-visual gradient describes the spatial variation in functional connectivity profiles across the cortex, highlighting the functional segregation between sensorimotor regions and the visual cortex (Larivière et al., 2020; Margulies et al., 2016). During the third trimester, this increasing segregation in gradient axis between the sensorimotor and visual regions may prioritize information communication within specific modality regions, which is essential for infants’ cognitive growth. Interestingly, the sensorimotor-to-visual gradient predicts the cognitive scores instead of motor and language at 2 years of age. A plausible explanation is that the establishment of the sensorimotor-to-visual gradient not only enables the sensory-specific functions at birth (Buckner & Krienen, 2013), but also the global efficient integration of multimodal information, which supports the maturation of infants’ cognitive abilities. Moreover, the anterior-to-posterior gradient, which were already evident at around 34 postmenstrual weeks, was not found to be associated with the cognitive growth in the first 2 years. This gradient, which is widely observed in the mammalian brain, could be related to the temporal differences in neurogenesis, reflecting the homology of the human brain (Cahalane et al., 2012; Elston, 2000; Rakic, 2002).

### Limitation

Several issues need to be considered. Firstly, previous research has indicated rapid changes in the structural connectome of the human brain during the preterm period (Zhao et al., 2019), and the sensorimotor-to-visual gradient has been closely linked to cortical morphology and white matter microstructures in term-born brains (Larivière et al., 2020). Our current study delineates the developmental process of core gradients in functional connectome during the third trimester. However, the relationship between the development of functional gradients and structural organization prior to birth still requires further investigation. Secondly, preterm neonates, compared to term-born infants, experience premature exposure to the external environment and are at higher risk of abnormal neurodevelopment of functional organization (Ball et al., 2013; Eyre et al., 2021). Future studies utilizing advancing fetal imaging technologies may help mitigate the impact of preterm birth on the development of brain functional gradients, thereby providing a more comprehensive understanding of longitudinal developmental trajectories in utero (Studholme, 2011). Finally, given the limited sample size and analytical techniques used in this study, further validation is necessary to confirm the associations between the functional gradients at birth and neurocognitive performance at 2 years of age. Future studies incorporating larger sample sizes and advanced methodologies will contribute to stronger evidence in this regard.

## STAR★Methods

### Participants

We collected resting-state fMRI data from 52 neonates recruited from Parkland Memorial Hospital in Dallas. A neonatologist and an experienced pediatric neuroradiologist screened the neonatal ultrasound and clinical MRI data as well as the neonatal and mothers’ medical records. Specific exclusion criteria were introduced in our previous studies (Xu et al., 2019; Zhao et al., 2019). Besides, we also performed a quality control for brain imaging data and then excluded 13 neonates considering their excessive head motion. Finally, we performed further analysis using data from 39 neonates (28 males/11 females; postmenstrual age at birth: 33.2 ± 4.5 weeks, range 25.1–40.7 weeks; postmenstrual age at scan: 37.0 ± 2.7 weeks, range 31.3–41.7 weeks). For each neonate, we obtained written and informed parental consents from neonatal parents. And this study was approved by the Institutional Review Board of the University of Texas Southwestern Medical Center.

### Follow-up neurodevelopmental assessments at 2 years of age

Among the 39 neonates used in further analyses, we only collected the follow-up neurodevelopmental assessments from 26 neonates (19 males/7 females, postmenstrual age at scan: 36.7 ± 2.8 weeks) at age of 2 years (chronological age at assessment: 23.5 ± 2.3 months). We assessed individual neurodevelopment using Bayley Scales of Infant and Toddler Development-third edition, including cognitive component scale, language component scale, and motor component scale (Bayley, 2006). The Bayley Scale III is a widely used standardized tool to assess the development of multiple dimensional functions of infants and toddlers. Specifically, the cognitive component scale assesses the non-verbal general cognitive ability, including exploration and manipulation, object relatedness, concept formation, and memory; the language component scale estimates the abilities including receptive and expressive communication through words and gestures; the motor component scale measures the development of gross and fine motor (Bayley, 2006). On the other hand, to avoid the Hawthorne effect, we asked a certified neurodevelopmental psychologist, who was unknown of the clinical conditions of neonates, to conduct the assessments for all toddlers.

### Image Acquisition and Preprocessing

All neonates were scanned with identical protocols using a single scanner. After neonates were naturally sleep, high-resolution T2-weighted (T2w) images and rs-fMRI data for each neonate were acquired using a Philips 3.0 T Achieva MR scanner at the Children’s Medical Center, Dallas. During the whole scanning period, using the earplugs, earphones, and extra foam padding to reduce the noise. Parameters for the T2w scans were as follows: repetition time (TR) = 3000 ms; echo time (TE) = 80 ms; field of view (FOV) = 168 × 168 mm^2^; number of slices = 65 and voxel size = 1.5 × 1.5 × 1.6 mm^3^. The rs-fMRI scans were performed using a T2w gradient-echo EPI sequence with the following parameters: TR = 1500 ms; TE = 27 ms; flip angle = 80°; FOV =168 × 168 mm^2^; number of slices = 30; number of volumes = 210; and voxel size = 2.4 × 2.4 × 3 mm^3^

All rs-fMRI data was preprocessed using SPM12 (Welcome Department of Cognitive Neurology, London, UK; www.fil.ion.ucl.ac.uk/spm), the Data Processing & Analysis for (Resting-State) Brain Imaging (DPABI) toolbox 4.3 version (Yan et al., 2016), and GRETNA 2.0 version (Wang et al., 2015) implemented in the MATLAB 2018b (Math Works, Natick, MA) platform.

The preprocessing of the rs-fMRI data included the following steps: (*i*) removal of the first 15 time points; (*ii*) slice timing correction; (*iii*) head motion correction and exclusion of neonates with excessive head movement (maximum head motion > 5 mm or > 5°, or mean frame-wise displacement > 1 mm); (*iv*) normalization of functional images to the customized group-level template. Specifically, we first normalized the functional images to individual T2w structural images, and then registered the individual T2w images to a publicly 37-week brain template (Serag et al., 2012). Then we re-registered individual T2w images using the group averaged first registered T2w image as the group-level template. Finally, the functional images were re-normalized to the group-level template using the transformation parameters from the second registration and resampled to a 3 mm isotropic resolution; (*v*) smoothing of the time series signal (FWHM = 4 mm); (*vi*) detrending of linear drift; and (*vii*) removal of spurious variance through linear regression. Nuisance regressors included Friston’s 24 head motion parameters, white matter, cerebrospinal fluid signals, and global brain signals; (*viii*) temporal bandpass filtering (0.01-0.08Hz); and (*ix*) scrubbing voxel-specific head motion by linear regression bad time point (FD>0.2 mm), and exclusion of neonates with excessive bad time points (> 50%) (Yan et al., 2013).

### Connectome Gradient Analysis

For each neonate, we performed connectome gradient analyses followed the method proposed by Margulies et al (Margulies et al., 2016). In brief, we first obtained a voxel-wise functional connectivity matrix (6,096 × 6,096) for each individual by calculating the Pearson correlation coefficient between the BOLD time series of each pair of gray matter nodes (without sub-cortical tissues). To avoid the noise of spurious connectivity, we thresholded the functional connectivity matrix of the top 10% correlation values of each voxel (Margulies et al., 2016). Then we calculated the cosine similarity between functional connectivity profiles of all pairs of voxels, and obtained a similarity matrix that represents the similarity in connectivity profiles of voxels. Next, we applied the diffusion map embedding technique (Coifman et al., 2005) to dimensionally reduce the high-dimensional similarity matrix to multiple low-dimensional embedding gradients followed previous parameters (Margulies et al., 2016; Y. Xia et al., 2022). To ensure the individual gradients were comparable, we used the Procrustes rotation approach (Langs et al., 2015) to align the original gradients of all neonates to the iterative template of term-born infants. Specifically, we first calculated the group level gradient of term-born neonates based on the group-averaged functional connectivity matrix, and then aligned raw gradients of all neonates to the group level gradient of term-born. The group-averaged aligned gradient of all neonates was taken as the gradient template for the second time alignment. We repeated this alignment process 100 times and obtained aligned gradients of each neonate. For the aligned individual gradients, we calculated the maturation index (i.e., the similarity with the gradient axis of the oldest age window) and the three global measures of the first two gradients, including explanation ratio (i.e., accounted for the variance of high-dimensional functional connectome), gradient range (i.e., difference between the values of positive and negative extremes of the gradient axis), and gradient variation (i.e., the standard deviation of values of gradient axis).

### General linear model

To investigate age-related changes in core functional connectivity gradients, we used the general linear model (Mardia et al., 2006) to assess age effect on global gradient measures and regional gradient scores of first two gradients in neonates, with gender and head motion parameters (mean FD) as covariates. The general linear model is defined as follows:

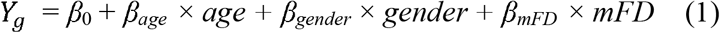

where *Y*_*g*_ is the connectome gradient measures of each neonate, *β*_*age*_ is the age effect. Bonferroni correction was performed to correct for multiple comparisons (8 times) across the global measures, and the Bonferroni corrected *P* < 0.05 was considered significant.

Further, to correct for multiple comparisons in the statistical analysis of regional gradient scores, we used a voxel-level threshold of *P* < 0.001 and a Gaussian Random Field (GRF) cluster-level correction of *P* < 0.05 to minimize false positive error.

### Prediction of the follow-up neurocognitive scores using support vector regression

To investigate whether the core functional connectome gradient in neonates can predict the follow-up neurocognitive maturation, we used the support vector regression (SVR) method with a linear kernel function to test the prediction power of functional gradients in neonates on follow-up neurocognitive scores at 2 years of age. Prediction performance was evaluated using the 5-fold cross-validation. Specifically, each training set includes 21 neonates and generated a classifier to predict the neurocognitive score of the remaining neonates. The classifier operated on the regional scores of the sensorimotor-to-visual gradient. To assess the prediction accuracies of each kind of neurocognitive score, the actual and predicted neurocognitive scores were compared using Pearson correlation analyses. The support vector regression analysis was implemented in the MATLAB2021b (Math Works, Natick, MA) platform using a modified *fitrsvm*() function. Finally, we used the permutation test (1,000 times) with randomly shuffled neurocognitive scores to examine whether the prediction performance exceeded the chance level.

## Acknowledgements

The work was supported by the National Key Research and Development Project (2018YFA0701402), the National Natural Science Foundation of China (31830034 and 82021004), and National Institute of Health (Grant Nos. R01MH092535, R01MH125333, R01EB031284, R01MH129981, R21MH123930, and P50HD105354).

## Data and materials availability

All statistical data and methods used in this study are available in the main text or supplementary materials.

